# Systematic investigation of the link between enzyme catalysis and cold adaptation

**DOI:** 10.1101/2021.08.25.457639

**Authors:** Catherine D. Stark, Teanna Bautista-Leung, Joanna Siegfried, Daniel Herschlag

**Affiliations:** Department of Biochemistry, Stanford University, Stanford, California 94305, United States; ChEM-H, Stanford University, Stanford, California 94305, United States; Department of Chemical Engineering, Stanford University, Stanford, California 94305, United States

## Abstract

Cold temperature is prevalent across the biosphere and slows the rates of chemical reactions. Increased catalysis has been predicted to be a general adaptive trait of enzymes to reduced temperature, and this expectation has informed physical models for enzyme catalysis and influenced bioprospecting strategies. To broadly test rate as an adaptive trait to cold, we paired kinetic constants of 2223 enzyme reactions with their organism’s optimal growth temperature (T_Growth_) and analyzed trends of rate as a function of T_Growth_. These data do not support a prevalent increase in rate in cold adaptation. In the model enzyme ketosteroid isomerase (KSI), there was prior evidence for temperature adaptation from a change in an active site residue that results in a tradeoff between activity and stability. Here, we found that little of the overall rate variation for 20 KSI variants was accounted for by T_Growth_. In contrast, and consistent with prior expectations, we observed a correlation between stability and T_Growth_ across 433 proteins. These results suggest that temperature exerts a weaker selection pressure on enzyme rate than stability and that evolutionary forces other than temperature are responsible for the majority of enzymatic rate variation.

## Introduction

Temperature is a ubiquitous environmental property and physical factor that affects the evolution of organisms and the properties and function of the molecules within them. As reaction rates are reduced at lower temperatures (Arrhenius, 1889; Wolfenden et al., 1999), the maintenance of enzyme rates has been suggested to be a universal challenge for organisms at colder temperatures that do not regulate their internal temperature (D’Amico et al., 2003; Fields et al., 2015; Siddiqui and Cavicchioli, 2006; Zecchinon et al., 2001). According to the rate compensation model of temperature adaptation, this challenge is met by cold-adapted enzyme variants providing more rate enhancement than the corresponding warm-adapted variants (Figure 1A). This model predicts that cold-adapted variants are faster than warm-adapted variants when assayed at a common temperature (Figure 1B). Indeed, this behavior has been reported for diverse enzymes, and these observations have been taken as support for this model (Figure 1C, Figure 1D) (Collins and Gerday, 2017; Feller and Gerday, 1997; Siddiqui and Cavicchioli, 2006; Smalås et al., 2000).

**Figure 1:**
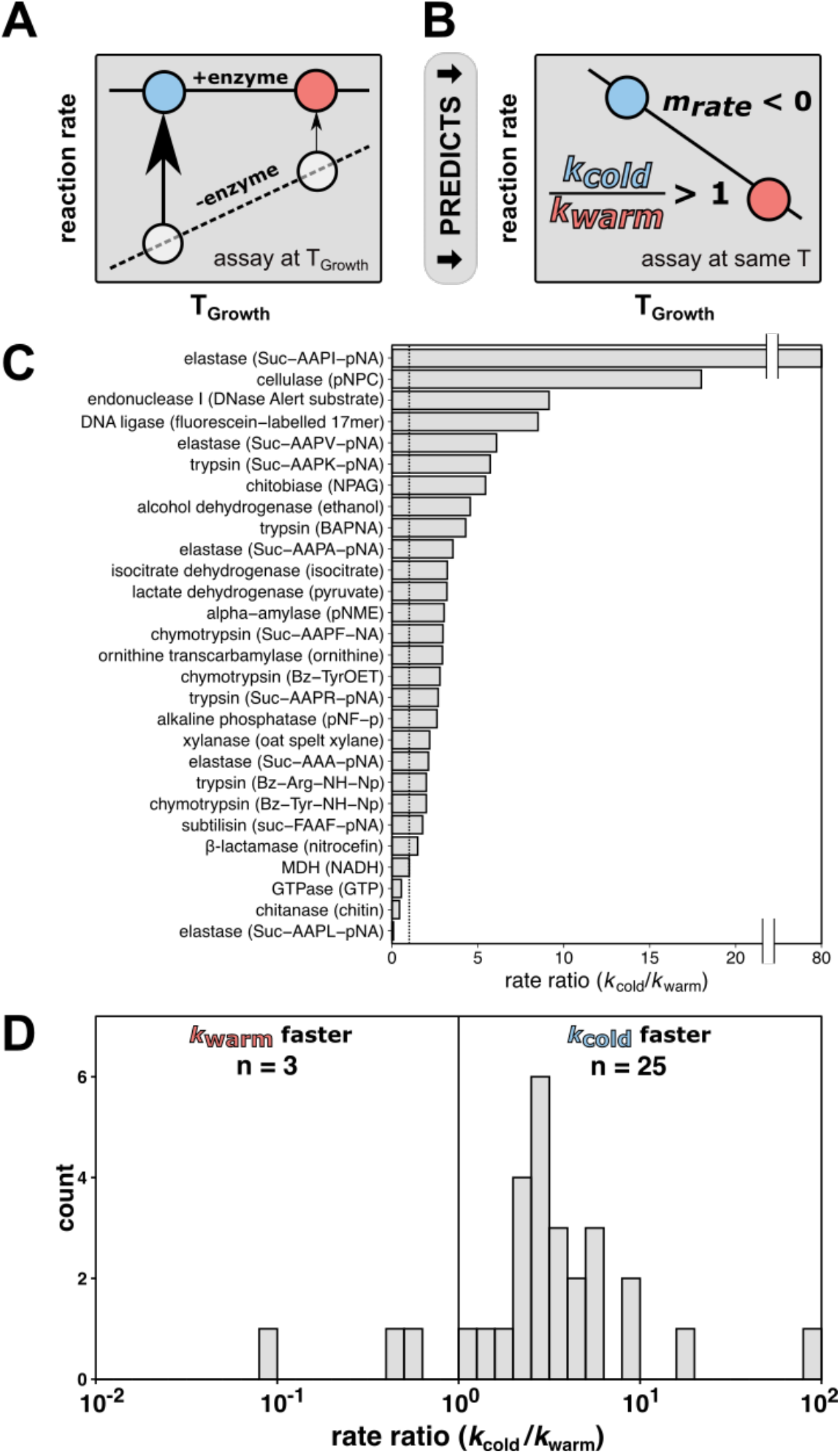
The rate compensation model of cold adaptation predicts that cold-adapted enzymes exhibit greater catalysis and are faster at a common temperature than their warm-adapted counterparts. (A) According to the rate compensation model of cold adaptation, a cold-adapted variant (blue circle) has larger rate enhancement than a warm-adapted variant (red circle). The dashed line represents the uncatalyzed reaction, the solid line represents the catalyzed reaction, and the arrows represent the rate enhancement at the respective organism T_Growth_. (B) When variants are assayed at a common temperature, rate compensation predicts a faster reaction for the enzyme from the cold-adapted organism, corresponding to a rate ratio (*k_cold_*/*k_warm_*) of greater than one and a negative slope of rate *vs*. T_Growth_ (*m_rate_*). (C, D) Rate comparisons of warm-adapted and cold-adapted enzyme variants made at identical temperatures from cold adaptation literature spanning indicated reactions with substrate specified in parentheses (Collins and Gerday, 2017; Feller and Gerday, 1997; Siddiqui and Cavicchioli, 2006; Smalås et al., 2000). The black vertical lines represents no rate change with temperature (*i.e.,* rate ratio = 1). **Figure 1—source data 1:** Figure1_SourceData1.csv

The observed rate effects (Figure 1C, Figure 1D) have also led to proposals of general physical models for cold adaptation linked to flexibility, as outlined in Supplementary file 1 (Åqvist et al., 2017; Arcus et al., 2016; Nguyen et al., 2017; Saavedra et al., 2018). Further, features identified in comparative structural analyses of cold and warm-adapted enzymes, such as fewer surface hydrogen bonds and salt bridges (Cai et al., 2018), have been suggested to increase flexibility and thus increase catalysis (Mandelman et al., 2019; H. J. Park et al., 2018; S.-H. Park et al., 2018). Correspondingly, the study of cold adaptation may have the potential to provide generalizable insights into physical properties of enzymes that make them better catalysts, a longstanding challenge in the field (Blow, 2000; Hammes et al., 2011; Kraut et al., 2003; Ringe and Petsko, 2008). From a practical perspective, the prediction of enhanced catalysis by cold-adapted enzymes has motivated low-temperature bioprospecting for biocatalysts to use in industrial processes (Bhatia et al., 2021; Bruno et al., 2019; Kuddus, 2018; Santiago et al., 2016).

Given the theoretical and practical implications of the proposed relationship between enzyme rate and organism growth temperature, we sought to test the generality of the rate compensation model of temperature adaptation. We collated enzyme rate data (Chang et al., 2021) and organism optimal growth temperature (T_Growth_) (Engqvist, 2018) for 2223 reactions using public databases. The results revealed no enrichment of faster reactions with colder growth temperatures, and thus did not support increased rate with decreasing environmental temperature as a prevalent adaptation in nature. Further, we found that most rate variation for the enzyme ketosteroid isomerase (KSI) is not accounted for by T_Growth_ despite strong prior evidence for temperature adaptation within its active site (Pinney et al., 2021). In contrast, a similar broad analysis revealed that stability correlates with T_Growth_, as expected. Our results suggest that temperature exerts a weaker selection pressure on enzyme rate than stability and that other evolutionary forces are responsible for most enzymatic rate variation.

## Results

### Broadly testing the rate compensation model

To investigate temperature adaptation of enzyme rate, we paired rate data from the BRENDA database (Chang et al., 2021) to organism growth temperatures. We simplified organism temperatures that may span changing conditions (Doblin and van Sebille, 2016) by matching the species name associated with the enzyme variant with the organism optimal growth temperature (T_Growth_) (Engqvist, 2018). Of 76,083 *k*_cat_ values in BRENDA, we found that 49,314 were for wild-type enzymes. Of these data, 16,543 values matched to microorganisms with known T_Growth_ values. We selected reactions in the database with variants from more than one organism, spanning 7086 *k*_cat_ values for 2223 reactions across 815 organisms with at least two variants per reaction (Figure 2A). These reactions spanned a temperature range of 1°C to 83°C (Figure 2B).

**Figure 2:**
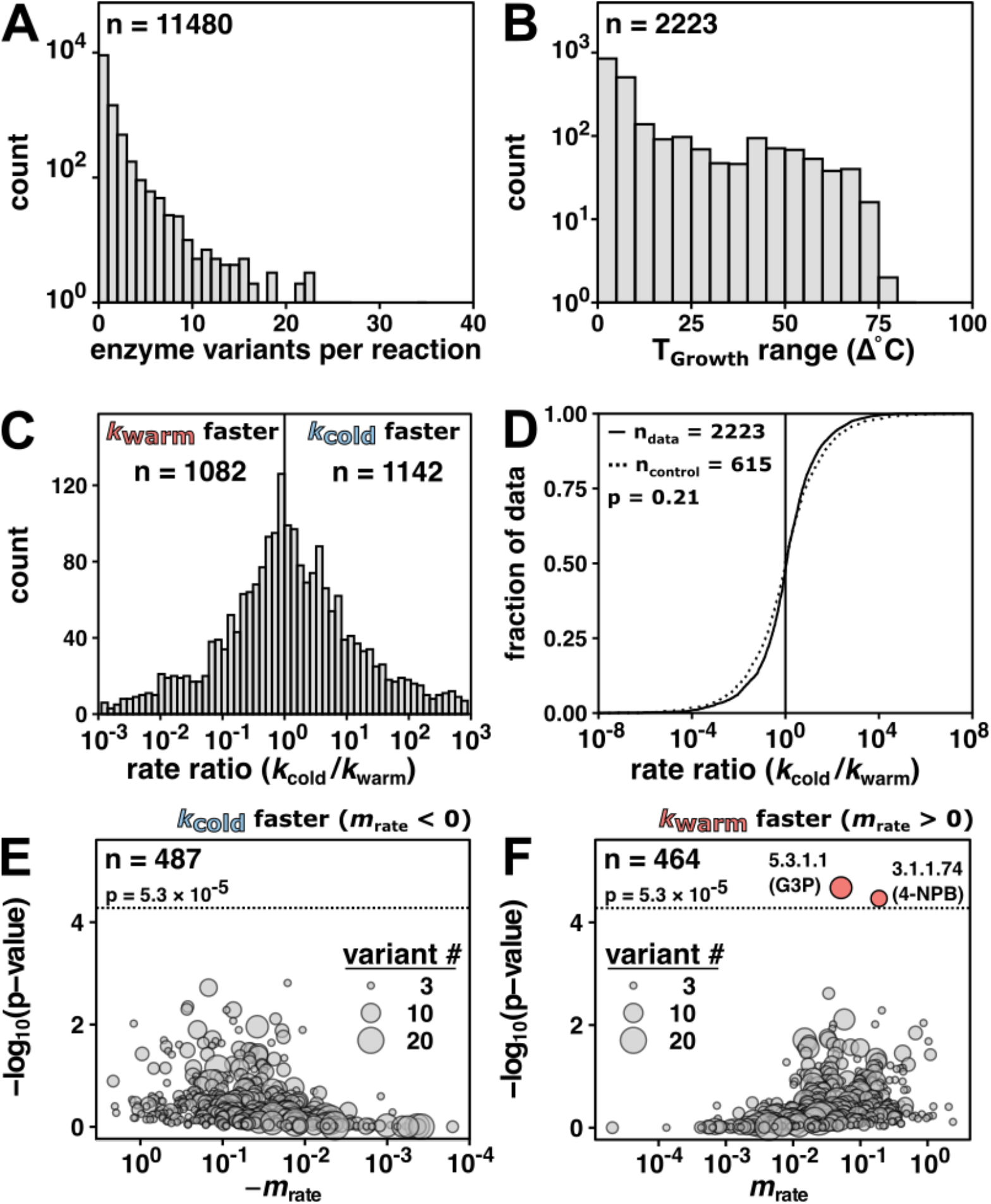
Enzyme rate data (*k*_cat_) do not indicate general rate compensation. (A) Enzyme variants per reaction of wild-type enzyme *k*_cat_ values (n = 11480 reactions) matched to T_Growth_. (B) Reactions with more than one enzyme variant (n = 2223 reactions). (C) Rate ratio distribution of the rate at the coldest T_Growth_ (*k_cold_*) divided by the rate of the variant from the warmest T_Growth_ (*k_warm_*) (median = 1.1 fold, 95% CI [1.00, 1.22], n = 2223 reactions). Vertical line at rate ratio = 1. For clarity, only data with rate ratios between 10^-3^ and 10^3^ are shown (>95% of the reactions). (D) Rate ratio (*k_cold_*/*k_warm_*) data (solid line, n = 2223 from panel C) compared to fold change control distribution (same T_Growth_; dashed line, median = 1.0 fold, 95% CI [0.89, 1.13], n = 615 reactions; p = 0.21, Mann–Whitney U test, two-sided). The black vertical line represents no rate change with temperature (*i.e.,* rate ratio = 1). (E, F) The significance and magnitude of the linear fit of reaction rate as a function of T_Growth_ for negative slopes (E, n = 487) and positive slopes (F, n = 464) in log space. E.C. number and (substrate) indicated for reactions significantly associated with temperature (Bonferroni correction; p-value < 5.3 × 10^-5^, n = 951). Dotted horizontal lines at p = -log_10_(5.3 × 10^-5^). 5.3.1.1: triose-phosphate isomerase; G3P: glyceraldehyde 3-phosphate; 3.1.1.74: cutinase; 4-NPB: 4-nitrophenyl butyrate. **Figure 2—source data 1:** Figure2_SourceData1.csv **Figure 2—source data 2:** Figure2_SourceData2.csv

For each enzyme reaction, we first calculated the rate ratio (*k_cold_*/*k_warm_*) between the rate of the variant from the lowest growth temperature organism and the rate of the variant from the highest growth temperature organism. We observed rate ratios greater than one (1142 reactions) as predicted by rate compensation, but nearly the same number of rate ratios of less than one (1082 reactions) (Figure 2C, *cf.* Figure 1D), providing no support for widespread or predominant rate compensation.

We also considered the distributions of rate ratios separated by assay temperature (25°C or 37°C; Figure 2—figure supplement 1A, 1B) and for wider T_Growth_ ranges (>Δ20°C or >Δ60°C; Figure 2—figure supplement 1C, 1D) to assess whether trends were obscured by mixed assay temperatures or narrow T_Growth_ ranges. However, no temperature-dependent trends emerged, supporting the above conclusion of an absence of widespread rate compensation.

To derive a control distribution, we compared enzyme variant rates originating from different organisms with identical T_Growth_ values. We found 615 reactions with more than one variant assigned the same T_Growth_, and we calculated the rate ratio and its reciprocal (*k_max_*/*k_min_* and *k_min_*/*k_max_*) for each reaction. This control distribution (dashed line, Figure 2D) was indistinguishable from the data distribution of rate ratios across T_Growth_ (solid line, Figure 2D; p = 0.21, Mann– Whitney U test, two-sided). Analogous analyses of *k*_cat_/*K*_M_ values lead to the same conclusions (Figure 2—figure supplement 2).

As it is not possible to prove the absence of a relationship (Altman and Bland, 1995), we examined the slope (*m*_rate_) of *k*_cat_ values *vs.* T_Growth_ for each of the 951 reactions with >2 variants (Figure 2D, Figure 2E, Figure 2—figure supplement 3) to address whether there might be a limited set of enzyme reactions exhibiting significant cold adaptation through a mechanism of enhanced rate. We found two reactions (triose-phosphate isomerase with glyceraldehyde 3-phosphate and cutinase with 4-nitrophenyl butyrate) significantly but positively associated with T_Growth_ (Bonferroni correction; p-value < 5.3 × 10^-5^, n = 951).

**Figure 3:**
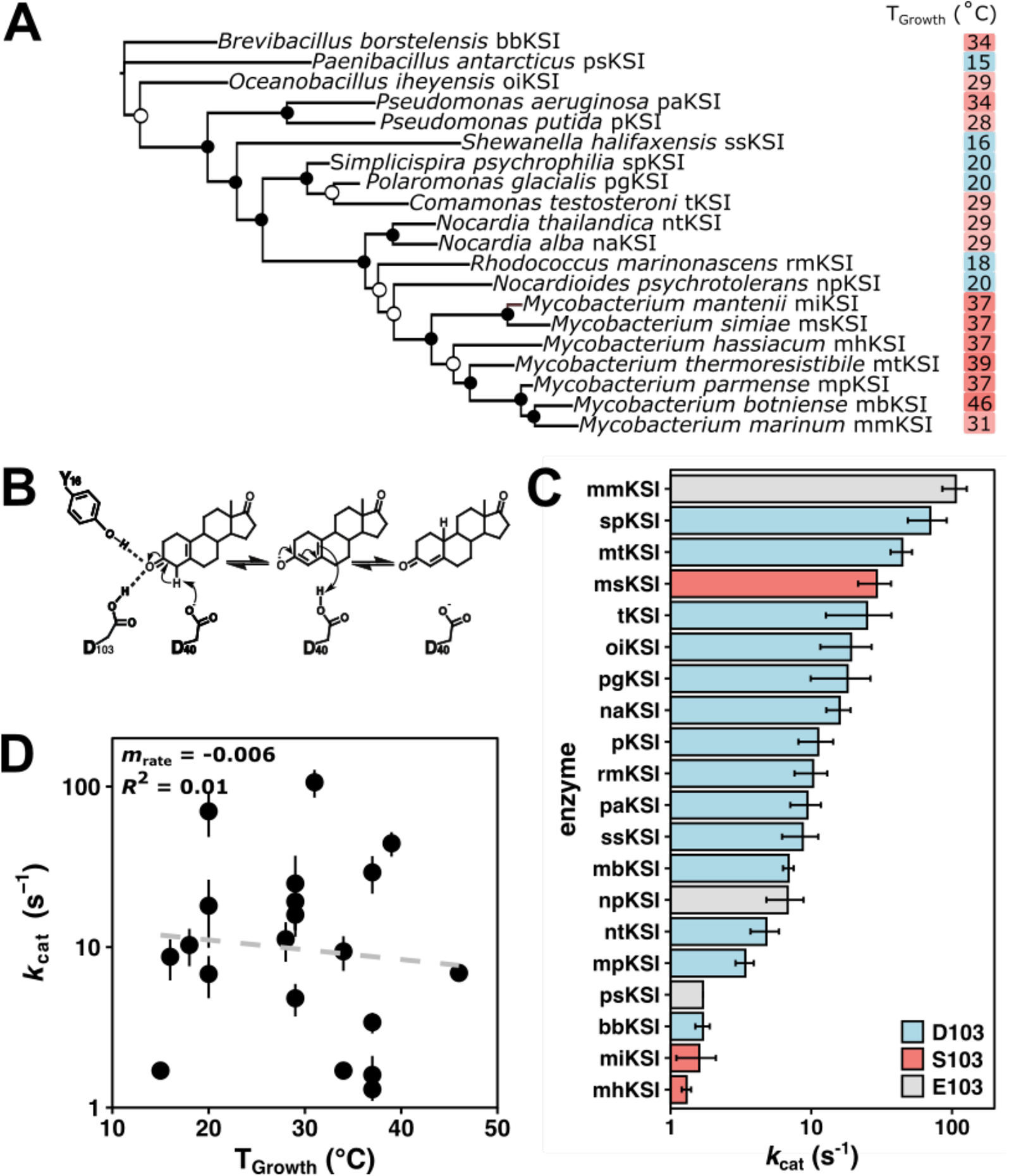
Ketosteroid isomerase (KSI) rates do not indicate rate compensation. (A) Unrooted maximum likelihood phylogenetic tree of KSI variants. Closed circles represent bootstrap values of >70%; open circles represent bootstrap values of 40-70%. (B) The mechanism of isomerization of the steroid 5(10)-estrene-3,17-dione by KSI. 5(10)-EST was used to allow direct measurement of the rate-limiting chemical step *k*_cat_ (Pollack et al., 1986) (C) Activity of KSI variants (*k*_cat_) at a common assay temperature of 25°C. Error bars represent standard deviation of at least two different experimental replicates varying [E] at least five-fold. KSI variants with D103 are represented in blue, S103 in red, and E103 in grey (*P. putida* numbering throughout). (D) Activity (log_10_(*k*_cat_)) of KSI variants at a common assay temperature (25°C) *vs.* organism growth temperature (T_Growth_) (n = 20, *m*rate = -0.006, *R*^2^ = 0.01, p = 0.02). **Figure 3—source data 1**: KSI origins and organism growth temperatures **Figure 3—source data 2**: Kinetic measurement of KSIs at 25°C with substrate 5(10)-estrene-3,17-dione. **Figure 3—source data 3**: Kinetic measurement of KSIs at 15°C with substrate 5(10)-estrene-3,17-dione.

In summary, the data provide no indication of rate increase as a consequence of decreasing T_Growth_. These results suggest that rate compensation is not a universal or prevalent consequence of temperature adaptation.

### Testing the rate compensation model for the enzyme ketosteroid isomerase (KSI)

To probe rate compensation in greater depth, we turned to the enzyme KSI for which recent data has demonstrated rate compensation (Pinney et al. 2021). Specifically, the change of a single active site residue at position 103 from serine (S103, prevalently found in warm-adapted KSI variants) to protonated aspartic acid (D103, prevalently found in mesophilic KSI variants) provided improved activity from a stronger hydrogen bond while also sacrificing stability by introducing an unfavorable protonation coupled to folding. We therefore used KSI to more deeply investigate the potential for rate compensation by assaying 20 variants that vary in sequence and T_Growth_ (Figure 3A).

KSI catalyzes the double bond isomerization of steroid substrates (Figure 3B) and is predicted to be part of a pathway that enables degradation of steroids for energy and carbon metabolism in bacteria (Horinouchi et al., 2010). KSI variants were identified by sequence relatedness to known KSIs. The 20 selected KSI variants ranged between 20–75% percent sequence identity to each other (Figure 3—figure supplement 1) and were selected from bacteria originating from environments spanning glaciers, ocean floor, soil, and wastewater with reported T_Growth_ values from 15°C to 46°C (Figure 3—source data 1). Each purified KSI demonstrated similar circular dichroism spectra at 5°C and 25°C, suggesting that variants were not unfolding at the 25°C assay temperature (Figure 3—figure supplement 2). All putative KSI variants exhibited isomerase activity on the steroid substrate 5(10)-estrene-3,17-dione (5(10)-EST) (Figure 3C).

We observed that the KSIs with the prevalent cold-adapted residue (D103 and the similar residue E103, *P. putida* numbering) were not uniformly faster than other KSIs in *k*_cat_ (Figure 3C) or *k*_cat_/*K*_M_ (Figure 3—figure supplement 3). The observation that one of the fastest variants contained serine at this position suggests that there are additional factors that influence its rate (Figure 3C & see Discussion).

For KSI, the value of *k*_cat_ decreased as a function of T_Growth_, but the shallow slope (*m*_rate_ = -0.006, p = 0.02) (Figure 3D) and the small coefficient of determination (*R^2^* = 0.01) of this relationship indicate that T_Growth_ accounts for little of the observed 80-fold rate variation. Similar activity trends were observed at an assay temperature of 15°C (Figure 3—figure supplement 3).

### Testing stability compensation using literature data

The absence of evidence for rate compensation led us to reinvestigate the widely accepted relationship between stability and growth temperature. Prior work has shown that temperature optima for observed enzyme rates correlate well with organism T_Growth_ (*r* = 0.75, (Engqvist, 2018)), but enzyme temperature optima reflect a combination of rate and stability effects. To isolate stability, we surveyed the relationship between stability and T_Growth_ using the ProThermDB, a collection of experimental data of protein and mutant stability (Nikam et al., 2021). Across 433 wild-type variants present in this database, we observed a significant relationship between T_m_ and T_Growth_ (Figure 4A, *R^2^* = 0.43, p = 2 x 10^-54^). For the 43 protein families with more than one reported variant, 39 had a higher melting temperature than their cold-adapted counterpart (Figure 4B).

**Figure 4:**
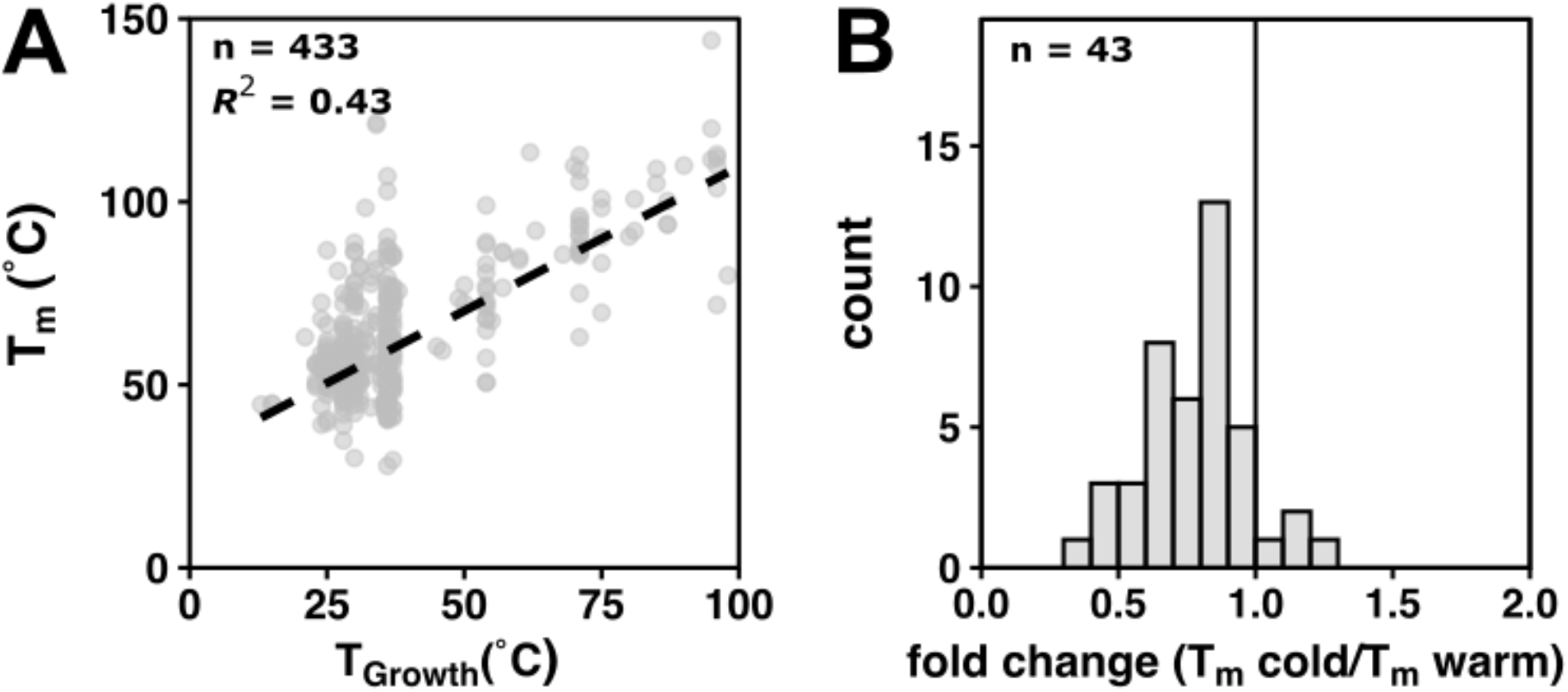
Protein stability data display stability compensation. (A) Wild-type T_m_ stability data from ProThermDB as a function of organism T_Growth_. Dashed black line represents a linear fit (n = 433, *R^2^* = 0.43). (B) Fold change (T_m_ cold/T_m_ warm) of wild-type protein variants (n = 43, median = 0.81, 95% [0.70, 0.85]. The black vertical line represents no change (*i.e.,* fold change = 1). **Figure 4—source data 1**: Figure4_SourceData1.csv **Figure 4—source data 2**: Figure4_SourceData2.csv

## Discussion

Enzymes have been widely posited to adapt to reduced temperature by increasing rate (Figure 1) (Collins and Gerday, 2017; D’Amico et al., 2003; Siddiqui and Cavicchioli, 2006; Zecchinon et al., 2001). Our results do not support this intuitive and common model as we found that cold-adapted enzyme variants are not generally faster than their warm-adapted counterparts. Even though there was prior evidence for temperature adaptation of the enzyme KSI that is accompanied by rate effects, we found that little of its overall rate variation was accounted for by organismal T_Growth_, suggesting instead that stability is the dominant driving force underlying the previously identified changes. Our observations suggest that enzyme rate is unlikely to be the primary trait selected for during adaptation to colder environmental temperatures, broadly and in the model system KSI.

Perhaps implicit in the expectation that catalysis will increase in cold adaption is the perspective that faster enzymes are better enzymes, with enzymes reacting at the diffusional limit denoted as “perfect” (Knowles and Albery, 1977). However, most enzymes operate well below the diffusional limit (Bar-Even et al., 2011), underscoring that an *optimal* reaction rate may be different than the *maximal* enzyme rate. There are multiple reasons why optimal or observed enzyme rates may differ from maximal rates. Rate optimization *in vivo* may be accomplished by altering gene expression (Somero, 2004), isoform expression (Somero, 1995), or cellular pH and osmolytes (Hochachka and Lewis, 1971; Yancey and Somero, 1979). Alternatively, the optimal enzyme rate may be lower than the maximal rate to channel metabolites and coordinate metabolism. Further, models of enzyme-metabolite pathway evolution predict that the subset of enzymes that govern pathway flux through rate-limiting steps are under strong rate selection (Noda-Garcia et al., 2018), and it is also possible that maximal enzyme rates are not evolutionarily accessible (Obolski et al., 2018). We speculate that rate compensation may be more probable for highly-related species that live in similar environments, such as marine species that live at different latitudes or depths but otherwise experience little environmental difference (Dong and Somero, 2009).

In contrast to our findings with rate, we observed strong evidence for stability compensation. The temperature dependence of protein unfolding (Becktel and Schellman, 1987) may exert a larger driving force on adaptation than the temperature dependence of rate. There may be an additional strong selection pressure to avoid unfolded states, as misfolded protein has been demonstrated to have deleterious fitness effects (Geiler-Samerotte et al., 2011) and cells expend considerable energy to clear misfolded variants using chaperones and degradation pathways (Clague and Urbé, 2010; Hartl et al., 2011; Lund, 2001). Additionally, adaptive paths towards stability may be more abundant and more accessible than analogous paths towards rate enhancement, given that each protein may be stabilized individually through a wide variety of mechanisms (Hart et al., 2014) and less constrained by biological context than an enzyme evolving synergistically with complex metabolic networks. The recent discovery of 158,184 positions from 1005 enzyme families that vary with growth temperature may further expand our understanding of the molecular strategies that underlie protein stabilization (Pinney et al., 2021).

The observation that one of the fastest KSIs contains the stabilizing but slowing active site residue, S103 (msKSI, Figure 3C), may illustrate some of the evolutionary complexity alluded to above. As observed with other KSIs, the S103D mutation in msKSI increases activity and decreases stability. However, in msKSI, the decreased stability from the S103D mutation renders it partially unfolded even in the absence of denaturants (Pinney et al., 2021). This result suggests a model where drift or other factors have led to an overall destabilized scaffold, such that msKSI cannot accommodate the activating S103D change (without unfolding) and has made other as yet unidentified amino acid changes that increase activity.

Flexibility has been posited to mechanistically link rate and stability, with multiple underlying interconnections discussed (see Supplementary file 1) (Åqvist et al., 2017; Arcus et al., 2016; D’Amico et al., 2003; Nguyen et al., 2017; Saavedra et al., 2018). Nevertheless, there are many degrees of freedom in an enzyme and most motions are not expected to be coupled to the enzyme reaction coordinate. Our observation of the absence of widespread rate compensation to temperature in contrast to observed stability compensation is consistent with this perspective, as are prior examples of enzyme stabilization in the absence of detrimental rate effects (Minges et al., 2020; Miyazaki et al., 2000; Siddiqui, 2017; Wintrode and Arnold, 2001; Zhao and Feng, 2018). A more complex relationship between these traits seems likely and underscores the need to relate individual and coupled atomic motions to overall flexibility, catalysis, and stability to unravel their intricate interconnections.

To understand why enzyme properties such as rate and stability measured with purified enzymes vary across organisms, we will need to determine their effects on fitness across biological and environmental contexts. Such studies may synergistically deepen our understanding of enzyme function, organismal evolution, and ecosystems.

## Acknowledgments

We thank M. Pinney, H. McShea, D. Mokhtari, C. Markin, I. N. Zheludev, J. Cofsky, E. E. Duffy and members of the Herschlag lab for thought-provoking discussions and review of this manuscript. We also thank F. Sunden, A. Chu, I. N. Zheludev, and F. Yabukarski experimental assistance and B. Eskildsen and D. Mokhtari for computational assistance. This research was supported by NSF Grant MCB-1714723, Stanford ChEM-H Chemistry-Biology Interface Training Program, and an NSF Graduate Research Fellowship to C.D.S and an NSF RET Supplement to J.S.

## Materials and Methods

### Literature Enzyme Rate Analysis

To capture enzyme rates reported throughout the literature, the BRENDA database was accessed using SOAP July 2021 (Chang et al., 2021) (www.brenda-enzymes.org) and the *k*_cat_ and *k*_cat_/*K*_M_ database entries retrieved by Enzyme Commission (E.C.) number were parsed for measurement value, substrate, rate, assay temperature, and variant (wild-type or mutant) status (Source Code 1). Microbial optimum growth temperature (Engqvist 2018) values from median organism optimal growth temperatures for microbes in culture (T_Growth_) were matched by organism name to rate entries. Rate data were filtered for *k*_cat_ and *k*_cat_/*K*_M_ values of wild type enzymes. Reactions are defined by E.C. number–substrate pair. The median value was taken in the case of multiple measurements of the same enzyme variant with the same substrate.

The rate ratio *k*_cold_/*k*_warm_ per reaction was determined by dividing rate of the enzyme from the organism with the minimum T_Growth_ by the rate of the enzyme from organism with the maximum T_Growth_. If a maximum or minimum T_Growth_ was shared between enzyme variants, then the median rate of the two variants was used in the rate ratio calculation. To account for enzyme rate variation arising independently of temperature, a control distribution from reactions with variants of the same T_Growth_ was derived. The fold change of the maximum value over the minimum value *k*_max_/*k*_min_ and its reciprocal *k*_min_/*k*_max_ was calculated for each reaction from the same T_Growth_ with at least two variants. To compare the rate ratio distribution of the data to the rate ratio control, the nonparametric two-sided Mann–Whitney U test was used with a significance threshold of p < 0.05. As no temperature-dependent trends emerged when data were restricted to measurements made at 25°C or 37°C or when the T_Growth_ range was limited to >Δ20°C and >Δ60°C, we used all data in the main analysis. We determined confidence intervals of the median parameters of the rate ratio distributions by bootstrap analysis (boot package in R, 10,000 replications) (Canty and Ripley, 2021; Davison and Hinkley, 1997). The *m*_rate_ values (slopes) per reaction were calculated by performing a linear regression relating the log_10_(rate) *vs*. organism T_Growth_. Significance threshold, corrected for multiple tests, was p < 5.33 × 10^−5^ (Bonferroni correction; p < 0.05/951).

### KSI Variant Identification, Cloning, Expression, and Purification

Putative ketosteroid isomerases (KSI) variants were identified by sequence relatedness to known KSI variants. Selection of variants was guided by associating putative KSI sequences with T_Growth_ by species (Engqvist, 2018). Seventeen variants were synthesized (GenScript or Twist Biosciences) and cloned (Gibson Assembly Protocol, New England Biolabs or Twist Biosciences) into pET-21(+) vectors. KSI variants were aligned using default parameters of Clustal Omega (Madeira et al., 2019) and the maximum likelihood tree was constructed using IQ-TREE with default parameters (Hoang et al., 2018; Nguyen et al., 2015). The constructs were expressed in *E. coli* BL21(DE3) cells and purified as previously described (Kraut et al., 2010).

### KSI Kinetic Measurements

The KSI substrate 5(10)-estrene-3,17-dione (5(10)EST) was purchased from Steraloids (Newport, RI). Reactions of purified KSIs with 5(10)EST were monitored continuously at 248 nm using a Perkin Elmer Lambda 25 UV/Vis spectrometer with an attached VWR digital temperature controlled circulating water bath (Pinney et al., 2021). Temperatures within the cuvettes were measured post-reaction using a platinum electrode thermistor (Omega Engineering) and the temperature of the circulating water bath was modified to maintain a constant internal cuvette temperature between reactions. Reactions were conducted in 40 mM potassium phosphate buffer, pH 7.2, 1 mM disodium EDTA, with 2% DMSO as a co-solvent to maintain substrate solubility. The kinetic parameters *k*_cat_ and *K*_M_ were determined by fitting the observed initial velocity of each reaction as a function of 5(10)EST concentration (9–600 µM; 6–7 different substrate concentrations per experiment) to the Michaelis–Menten equation. Reported values of *k*_cat_ and *K*_M_ are the average of 3–9 independent experiments with at least two different enzyme concentrations varied by at least 5-fold. Reported errors are the standard deviations of these values.

### KSI Circular Dichroism (CD)

CD spectra were collected for each KSI variant in 40 mM potassium phosphate buffer, pH 7.2, 1mM EDTA, at enzyme concentration 20 µM at 5°C and 25°C. Measurements were made on a J-815 Jasco Spectrophotometer between 190-250 nm at 1 nm bandwidth and 50 nm/min scanning speed in a 0.1 cm cuvette (Hellma Analytics).

### Literature Stability Analysis

Wild-type mutation type stability data were downloaded from ProThermDB (Nikam et al., 2021) with the following fields: protein information (entry, source, mutation, E.C. number), experimental conditions (pH, T, measure, method), thermodynamic parameters (T_m_, state, reversibility), and literature (PubMed ID, key words, reference, author). Wild type protein data were matched by species name to microbial optimal growth temperatures T_Growth_ (Engqvist, 2018).

---

**Figure 2—figure supplement 1:**
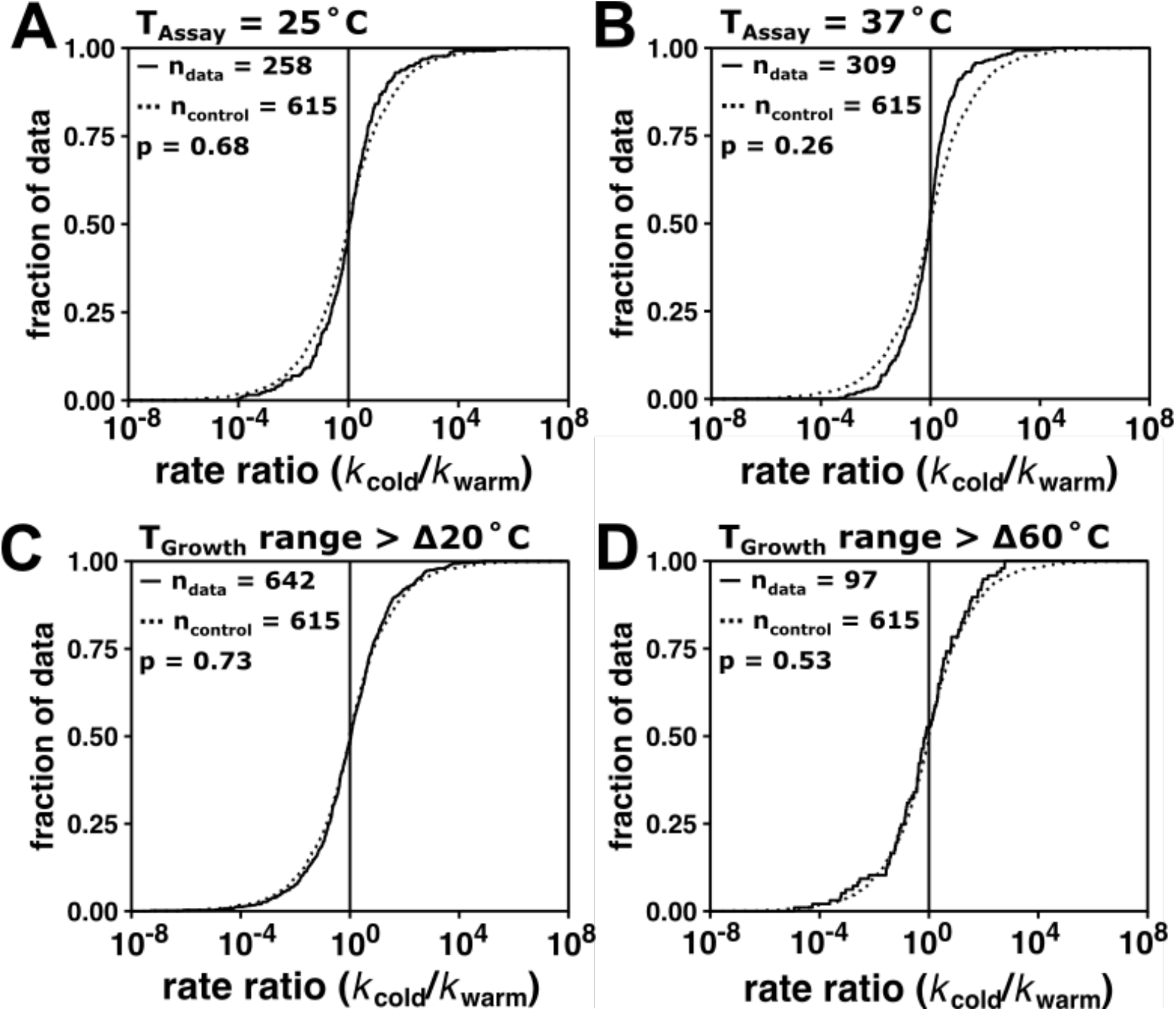
Specifying assay temperature and organism optimal growth temperature range per reaction does not alter conclusions. (A) Distribution of *k*_cat_ rate ratio values including only measurements made at 25°C and (B) 37°C. (C) Distribution of rate ratios with T_Growth_ range > Δ20°C and (D) T_Growth_ range > 60°C. Reported p-values from two-sided Mann– Whitney U test comparing filtered data (solid line) and the control data (dotted line, see Materials & Methods).

**Figure 2—figure supplement 2:**
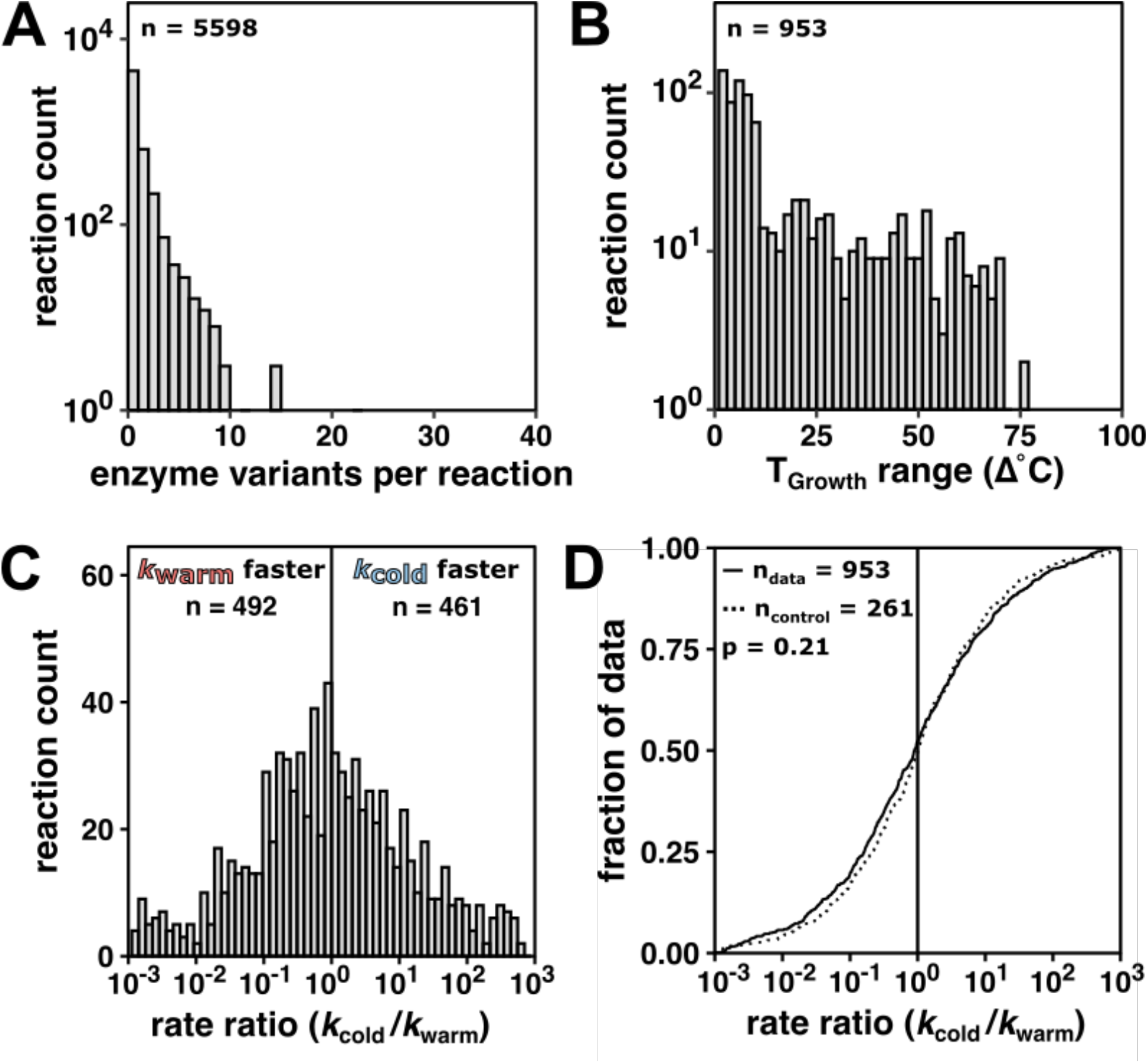
Enzyme rate data for *k*_cat_/*K*_M_ do not indicate rate compensation, supporting the conclusions from the *k*_cat_ analysis in the main text. (A) Variants per reaction of wild-type enzyme *k*_cat_ values (n = 5598 reactions) matched to T_Growth_. (B) Number of reactions spanning the specified T_Growth_ range (n = 953 reactions with >1 variant). (C) *k*_cat_/*K*_M_ rate ratio (*k*_cold_/*k*_warm_) distribution (median = 0.93 fold, 95% CI [0.78, 1.12], n = 953 reactions). Grey vertical line at rate ratio = 1. (D) *k*_cat_/*K*_M_ rate ratio (*k*_cold_/*k*_warm_) data (black line, n = 953 reactions) with *k*_cat_/*K*_M_ rate ratio control (grey line, median = 1.00 fold, 95% CI [0.82, 1.21], n = 307 reactions) determined in the same way as the *k*_cat_ rate ratio control in the main text (see Materials & Methods) (p = 0.80, Mann–Whitney U test, two-sided). For clarity, only data with rate ratios between 10^-3^ and 10^3^ are shown, representing >90% rate ratio data in (C) and >83% of rate ratio control values in (D). Black vertical line represents no rate change with temperature (*i.e.*, rate ratio = 1).

**Figure 2—figure supplement 3:**
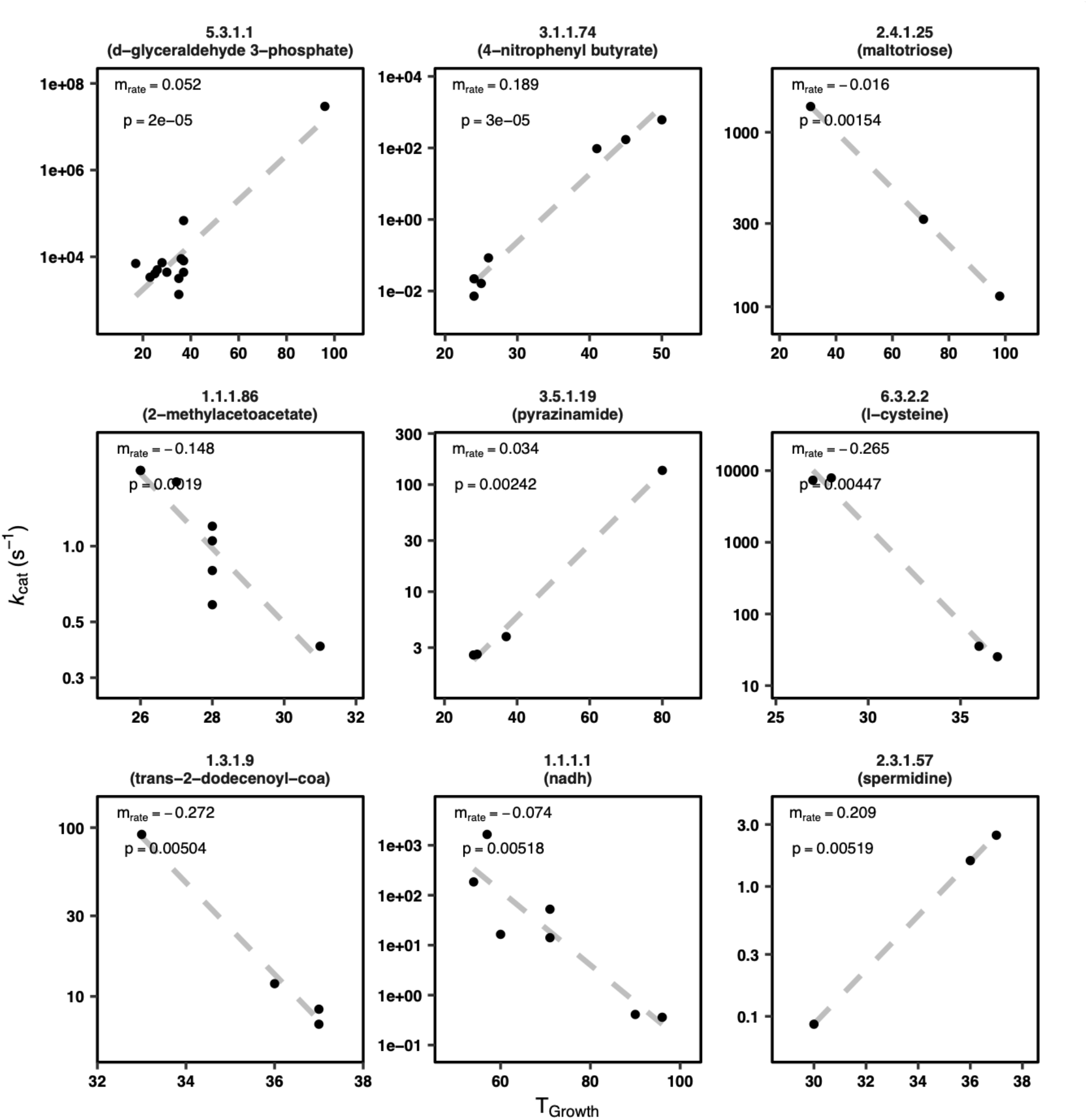
Example *m_rate_* plots (9 of 951 reactions shown). Reactions with the rate of constituent variants in order of *m*_rate_ p-value (for all reactions shown, p < 5.2 × 10^-3^). *m*_rate_ is the slope of log_10_(*k*_cat_) *vs.* T_Growth_. Note different scales for the axes. 5.3.1.1: triose-phosphate isomerase; 3.1.174: cutinase; 2.4.1.25: 4-alpha-glucanotransferase; 1.1.1.86: ketol-acid reductoisomerase; 3.5.1.19: nicotinamidase; 6.3.2.2: glutamate-cysteine ligase; 1.3.1.9: enoyl-[acyl-carrier-protein] reductase (NADH); 1.1.1.1: alcohol dehydrogenase; 2.3.1.57: diamine N-acetyltransferase.

**Figure 3—figure supplement 1:**
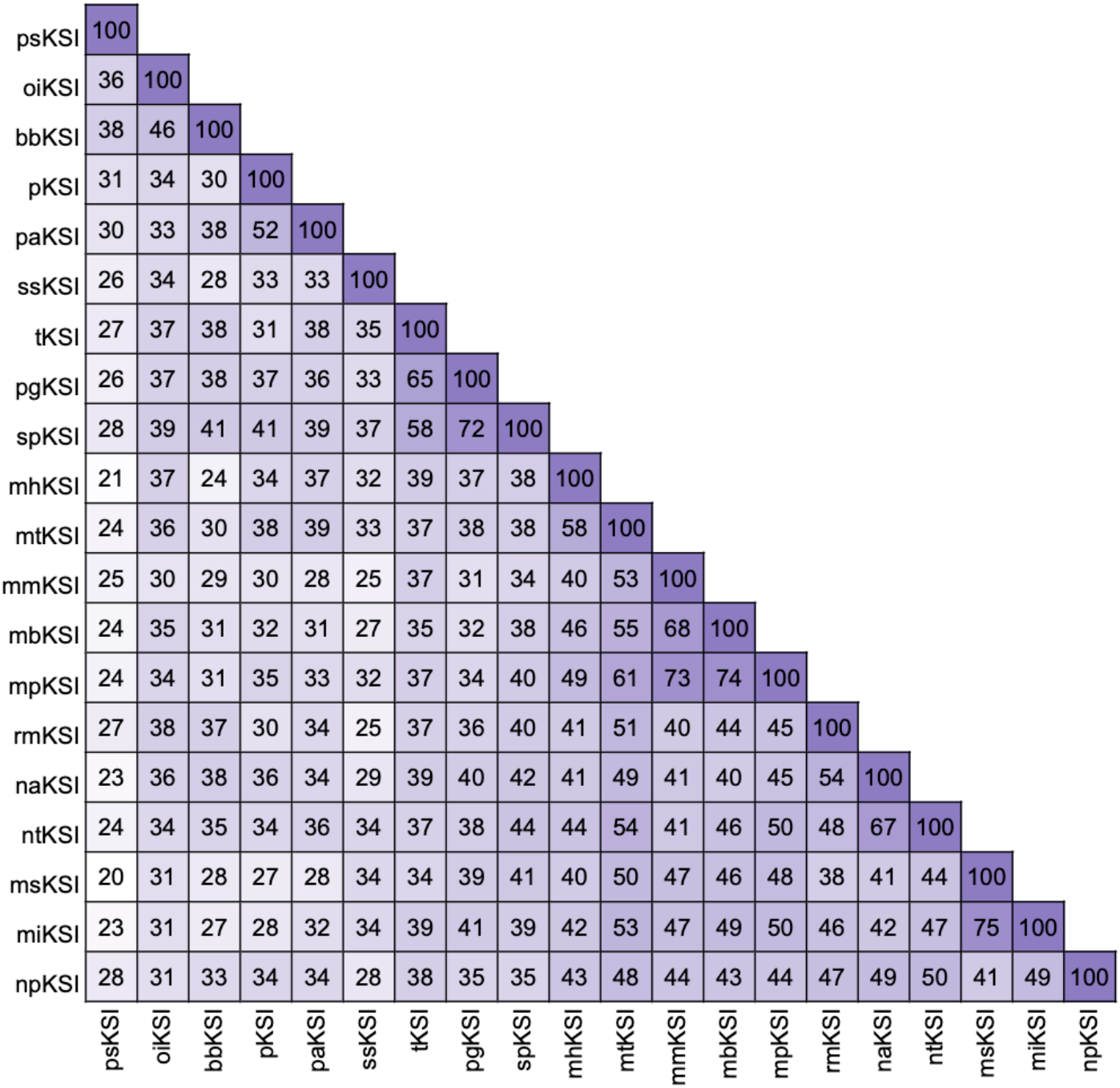
KSI variant similarity. The primary sequence variation of each KSI variant ranges from 20-75% amino acid identity.

**Figure 3—figure supplement 2:**
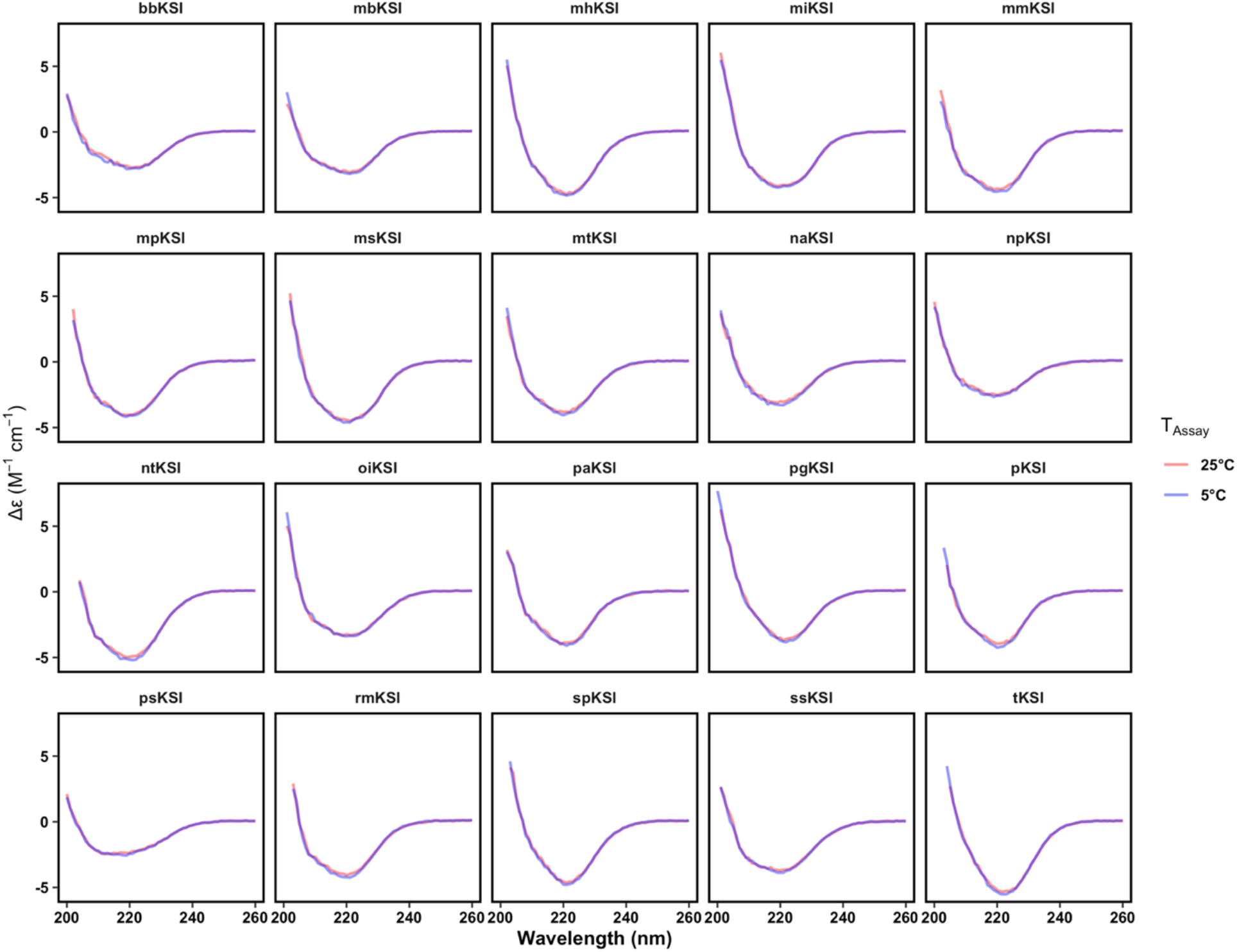
KSI variant circular dichroism spectra are similar at cold and warm temperature. Far ultraviolet circular dichroism (CD) spectra at 5°C (blue) and 25°C (red) are indistinguishable. Measurements for each variant were made at an enzyme concentration of 20 µM.

**Figure 3—figure supplement 3:**
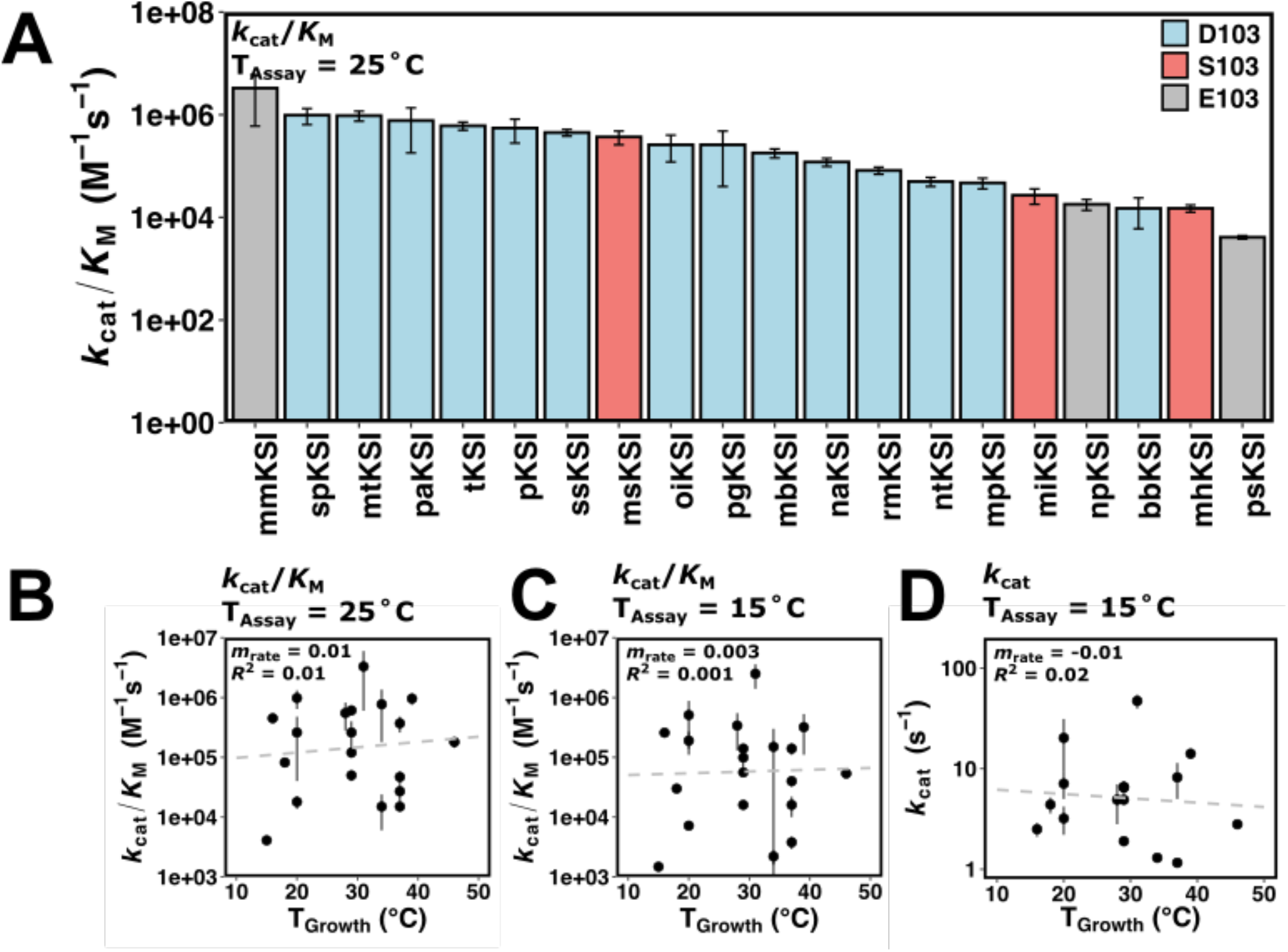
Ketosteroid isomerase rates vary with organism growth temperature in *k*_cat_ and in *k*_cat_/*K*_M_. (A) Rate of KSI variants (*k*_cat_/*K*_M_) at a common assay temperature (T_Assay_) of 25°C. KSI variants with D103 are represented in blue, S103 in red, and E103 in grey (*P. putida* numbering). (B) Rates (*k*_cat_/*K*_M_) of KSI variants at 25°C assay temperature (T_Assay_) *vs.* organism growth temperature (T_Growth_) (n = 20, *m*_rate_ = 0.01, *R*^2^ = 0.01, p = 4 ×10^-7^). (C) Rates (*k*_cat_/*K*_M_) of KSI variants at 15°C assay temperature (T_Assay_) *vs.* organism growth temperature (T_Growth_) (n = 20, *m*_rate_ = 0.003, *R*^2^ = 0.001, p = 3 ×10^-7^). (D) Rates (*k*_cat_) of KSI variants at 15°C assay temperature (T_Assay_) *vs.* organism growth temperature (T_Growth_) (n = 20, *m*_rate_ = -0.01, *R*^2^ = 0.02, p = -0.11). Error bars represent standard deviation of at least two different experimental measurements varying [E] at least five-fold (25°C) or two-fold (15°C).

**Figure 3—source data 1:**
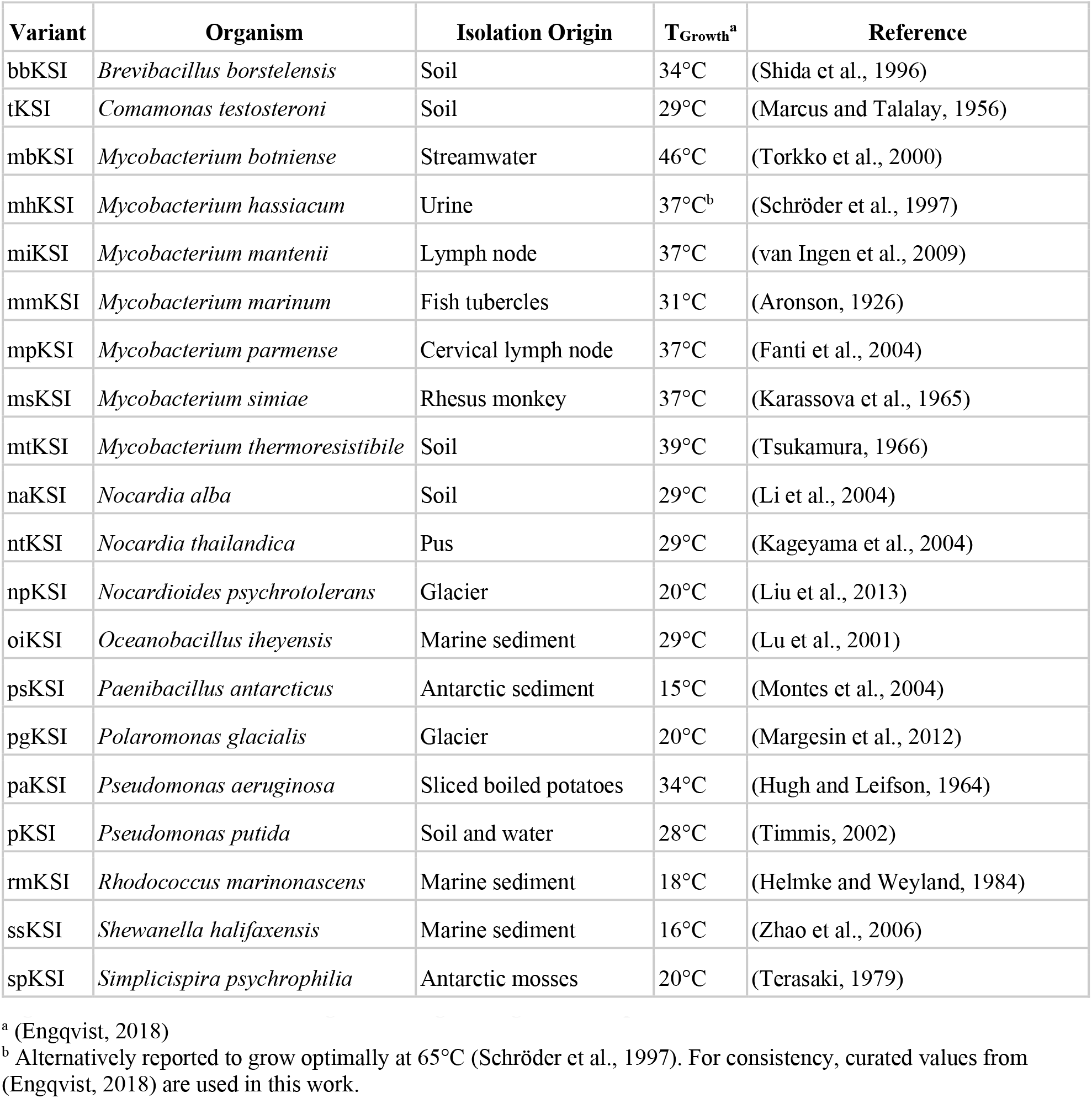
KSI origins and organism growth temperatures.

**Figure 3—source data 2:**
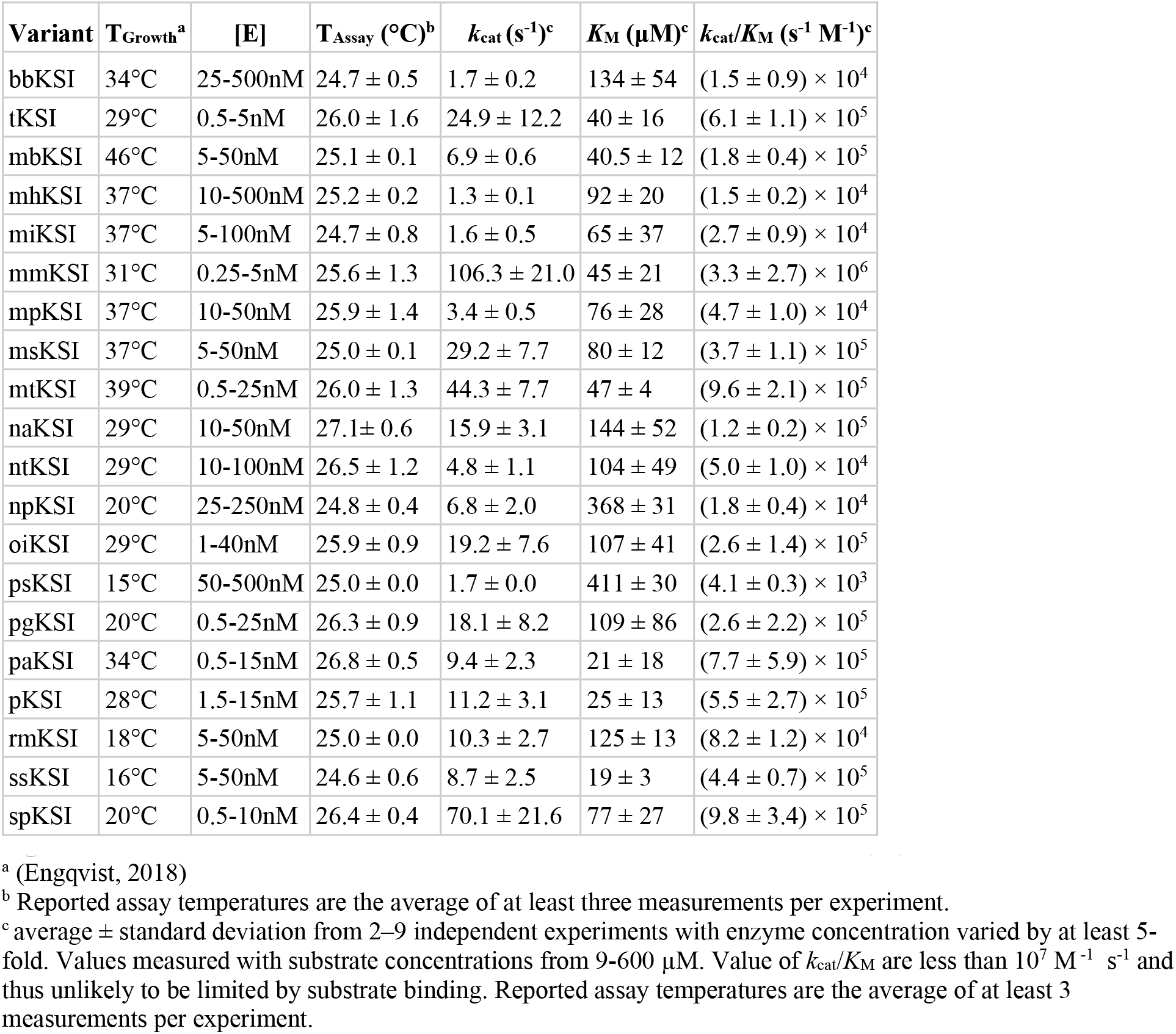
Kinetic measurement of KSIs at 25°C with substrate 5(10)-estrene-3,17-dione.

**Figure 3—source data 3:**
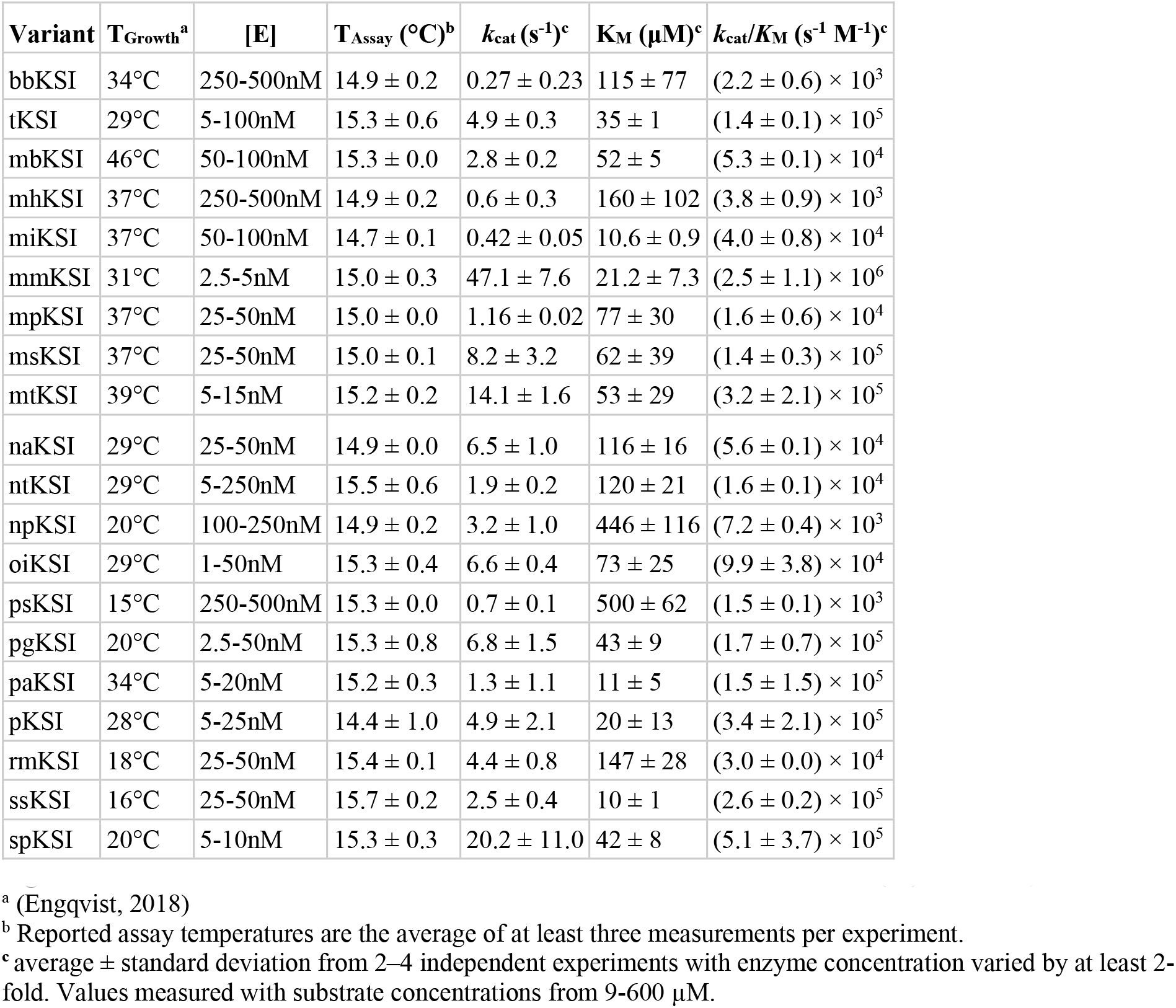
Kinetic measurement of KSIs at 15°C with substrate 5(10)-estrene-3,17-dione.

**Supplementary file 1.** Overview of proposed molecular models of cold adaptation.

| Trait | Model | Highlighted experimental observations | References |
| --- | --- | --- | --- |
| Flexibility (general) | Higher flexibility overcomes reduced motion at lower temperature, allowing enhanced catalysis. | Fish lactate dehydrogenase rate ($k_{cat}$) correlates with average body temperature. Mollusk cytosolic malate dehydrogenase substrate affinity ($K_M$) correlates with habitat temperature. Psychrophile α-amylase variant is faster than mesophilic and thermophilic variants. Chitobiase engineering stabilizes and reduces activity of psychrophilic enzyme. | (Feller and Gerday, 2003; Fields et al., 2015) |
| Flexibility (specific) | Surface flexibility decreases enthalpy and entropy activation terms, reducing temperature dependence of reaction. | Computational methods including molecular dynamics and empirical valence bond simulations of diverse enzyme systems, with specific focus on citrate synthase and trypsin variants, suggest importance of flexibility of surface residues in cold adaptation. | (Åqvist et al., 2017) |
| Heat capacity modulation | Altered temperature dependence of reaction reduces rate decrease as temperature is lowered | More negative $\Delta C_p^\ddagger$ has been observed in psychrophilic isopropylmalate dehydrogenase and α-glucosidase. In contrast, more negative $\Delta C_p^\ddagger$ has also been suggested as a driver of adaptation to higher temperature in reconstructed ancestral adenylate kinase sequences. | (Arcus et al., 2016; Nguyen et al., 2017) |
| Dynamic allostery | Partial unfolding arising from conformational entropy-enhancing mutation can affect $k_{cat}$ and $K_M$ | Dynamics-based regulation arising from mutations distal to the active site of mesophilic adenylate kinase affect substrate affinity and turnover, suggesting a mechanism of cold adaptation. | (Saavedra et al., 2018) |

**Supplementary File 2.**
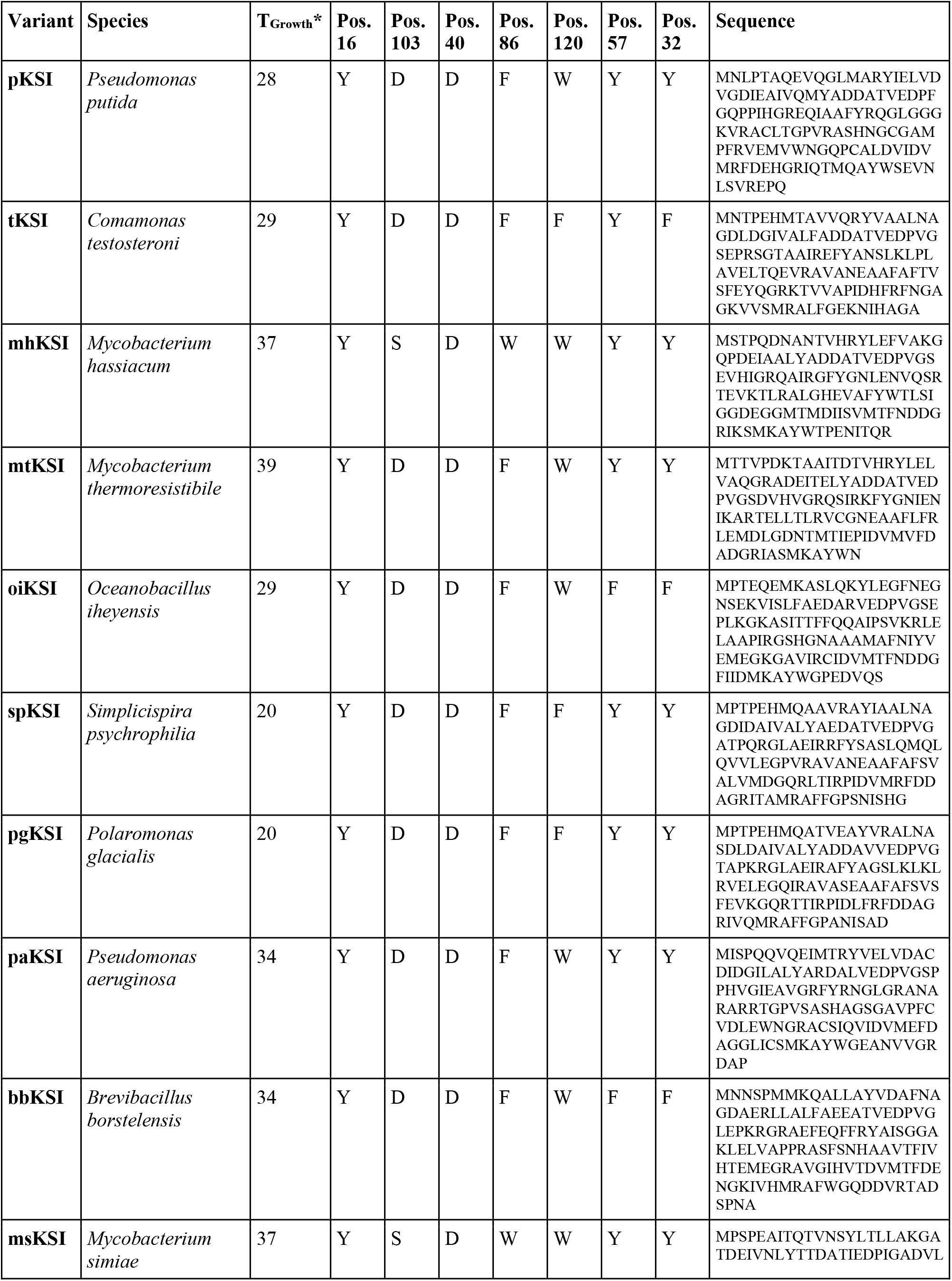
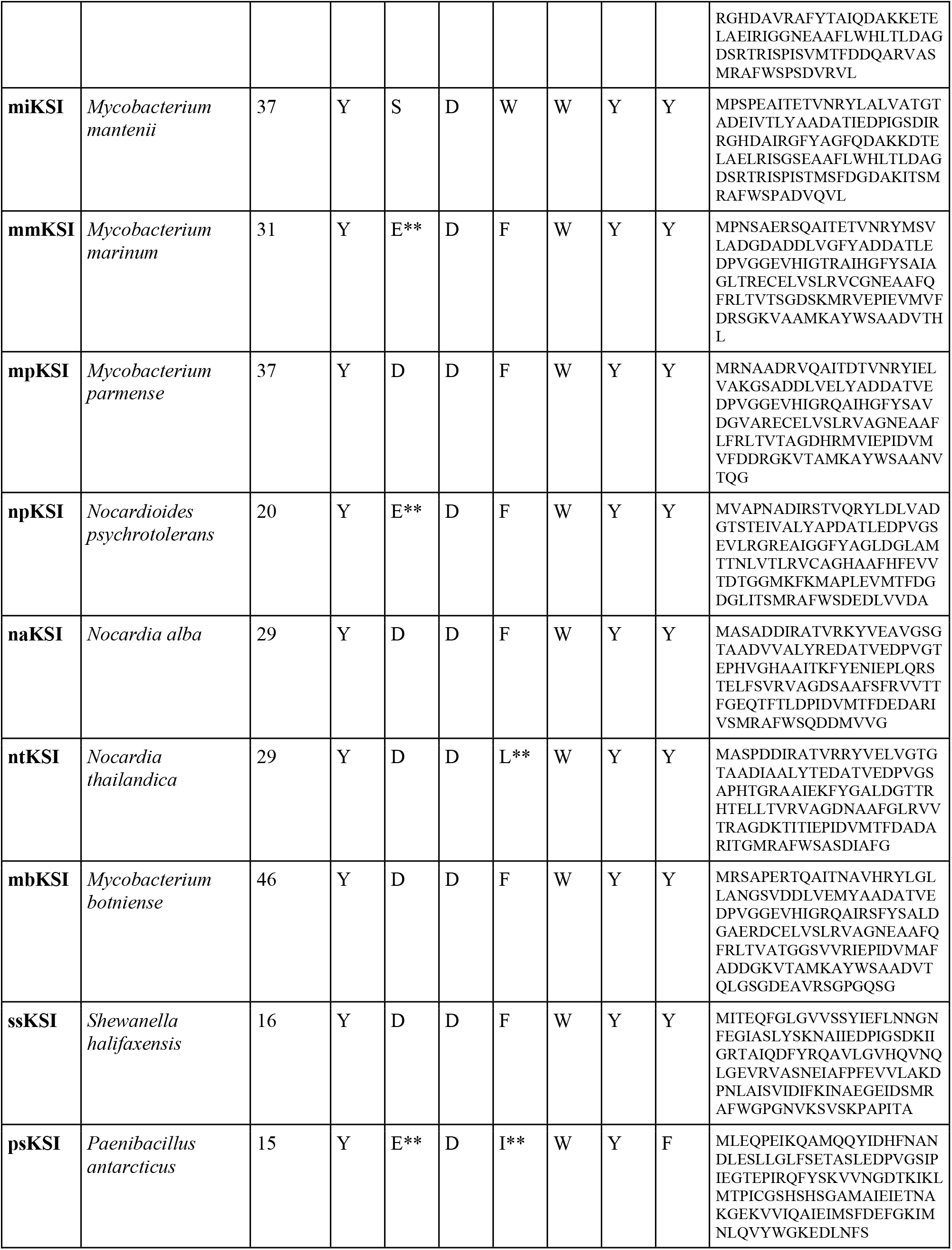
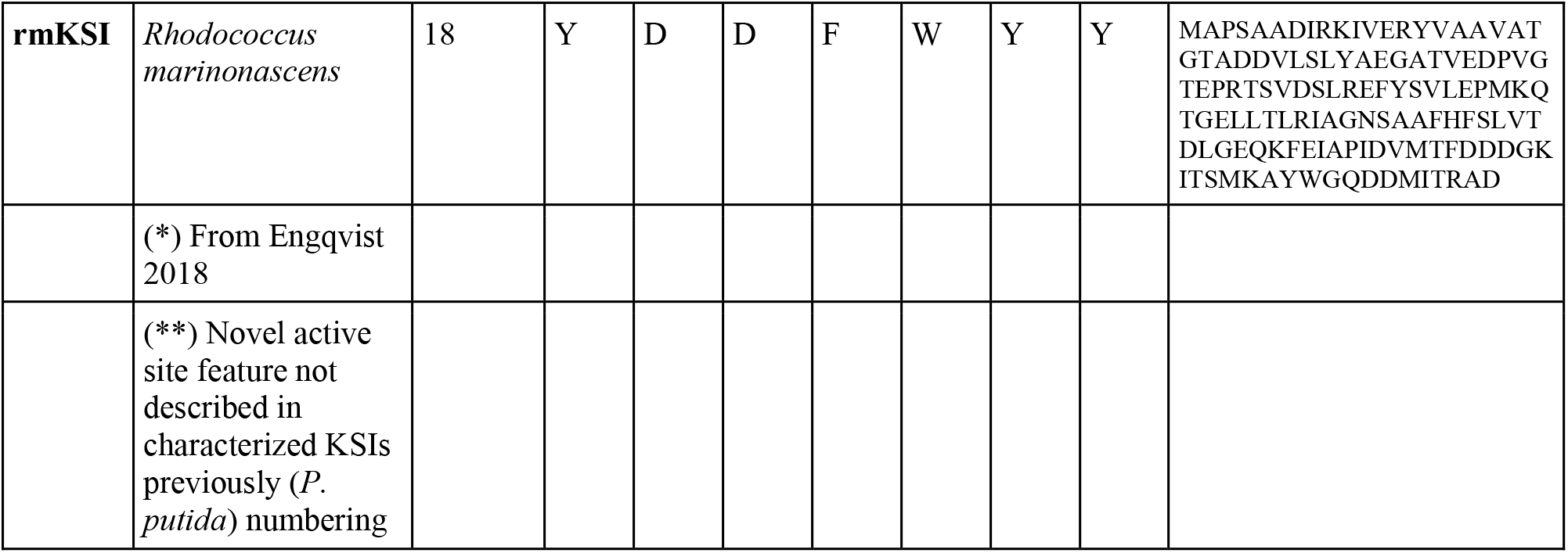
KSI sequences.

